## Supplementary material for "Cross-species Comparison Reveals Therapeutic Vulnerabilities Halting Glioblastoma Progression": Mathematical_modeling_supplement

### 1 Mathematical model

The current mathematical model departs from those previously introduced for healthy adult neurogenesis in the dentate gyrus of the hippocampus [3, 4] and the ventricular-subventricular zone (V-SVZ) [5]. The healthy V-SVZ model assumes a population of neural stem cells (NSC) that can reside in a quiescent state (qNSC) or activate and enter the cell cycle (aNSC). An aNSC divides into two daughter cells that can either be stem cells themselves (symmetric self-renewal) or more differentiated progeny called transient amplifying progenitors (TAP, symmetric differentiation). TAPs perform multiple rounds of division (amplification) and eventually divide into two neuroblasts (NB) that finally migrate through the rostral migratory stream to the olfactory bulb, thus leaving the system under consideration (see [6] for a review).

The model consists of a system of ordinary differential equations that encompass the transitions between the lineage cell types described above (1). The parameter set consists of $r$ - the activation rate of qNSCs into aNSCs; $b$ - the fraction of self-renewal, which represents the fraction of daughter cells that are NSCs themselves, and can be viewed as the probability that an aNSC divides into two qNSCs, as opposed to two TAPs; $p_A$ - the proliferation rate of aNSCs; $p_T$ - the proliferation rate of TAPs; and $\delta$ - the rate at which NBs leave the system considered. Additionally, $n$ represents the average number of divisions that a TAP can perform. In previous works of adult neurogenesis $n$ was assumed to be equal to 2 or 3 [3, 4, 5].

$$\begin{aligned}
\frac{dQ}{dt} &= -rQ(t) + 2bp_A A(t) \\
\frac{dA}{dt} &= rQ(t) - p_A A(t) \\
\frac{dT_0}{dt} &= 2(1-b)p_A A(t) - p_T T_0(t) \\
\frac{dT_i}{dt} &= 2p_T T_{i-1}(t) - p_T T_i(t) \\
\frac{dT_n}{dt} &= 2p_T T_{n-1}(t) - p_T T_n(t) \\
\frac{dN}{dt} &= 2p_T T_n(t) - \delta N(t).
\end{aligned} \tag{1}$$

Focusing on the study of GBM, as there exists no evidence of a possible maximum number of divisions $n$, we reduce the system to four ODEs describing the populations $Q$ (quiescent stem cells); $A_1$ (active stem cells) and $A_2$ (progenitor cells) which together form the $A$ population considered throughout the main manuscript; and $D$ (differentiated cells). As before, $r$ represents the activation rate of $Q$ into $A_1$. The fraction of self-renewal parameter $b$ is now denoted by $b_1$ representing the probability of an $A_1$ to give rise to two $Q$. Additionally, the set of TAP equations ($T_i$ compartments) are now collapsed into one ODE by introducing an additional parameter $b_2$ that represents the fraction of self-renewal of progenitors $A_2$. This is referred to as amplification in the main text. The crucial difference between $A_1$ and $A_2$ is that, whilst $A_1$ can give rise to more stem cells $Q$ that can activate and generate $A_2$ daughter cells, $A_2$ are more differentiated and can only give

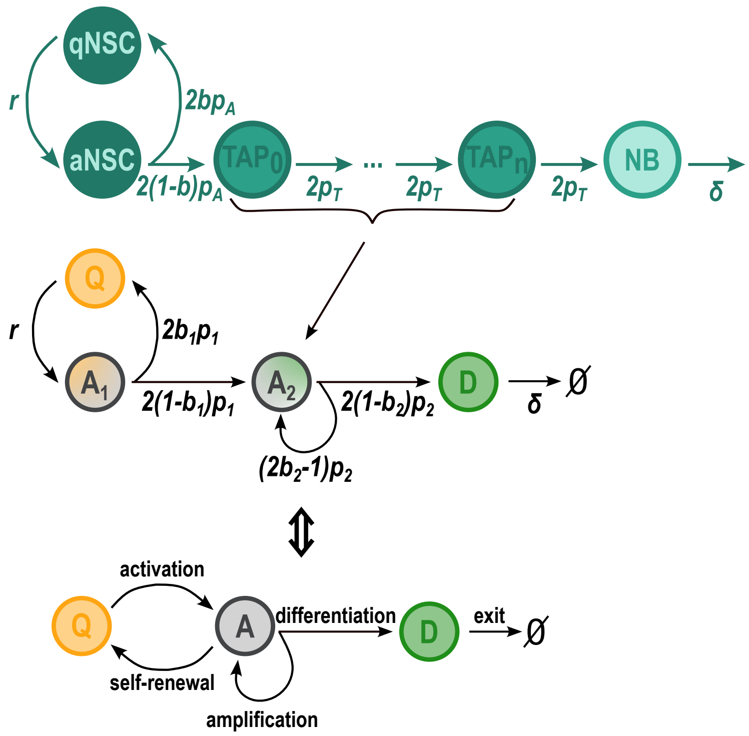


Figure 1: **S**chematics: Compartments of healthy neural cell types and their transitions represented in the mathematical model (1), collapsed into four compartments of GBM cell types given by the model (2, which describes the Q-A-D populations studied in the main manuscript.

rise to more $A_2$ or $D$. Finally, their proliferation rates are denoted $p_1$ and $p_2$, respectively and $\delta$ describes the exit of $D$ from the system, for example by death or migration, as before. As a result, the reduced system of ODEs describing GBM dynamics reads as in Eq. (2).

$$\begin{aligned}
\frac{dQ}{dt} &= -rQ(t) + 2b_1p_1A_1(t) \\
\frac{dA_1}{dt} &= rQ(t) - p_1A_1(t) \\
\frac{dA_2}{dt} &= 2(1-b_1)p_1A_1(t) - (2b_2-1)p_2A_2(t) \\
\frac{dN}{dt} &= 2(1-b_2)p_2A_2(t) - \delta D(t).
\end{aligned} \tag{2}$$

The QAD proportions considered in the main text are then described by the terms $Q/(Q+A_1+A_2+D), (A_1+A_2)/(Q+A_1+A_2+D), D/(Q+A_1+A_2+D)$, respectively. For clarity, we will denote here $Q_p, A_p, D_p$ for the respective QAD proportions.

#### 2 Stationary distributions of different cell types

Of note, mathematical analysis proves that the QAD proportions $(Q_p, A_p, D_p)$, converge to steady states. This is a property of linear systems of ODEs describing structured populations, governed by Metzler matrices (i.e., matrices that have non-negative values off the diagonal), which can be proved by employing the Peron-Frobenius theorem [1, 2]. Indeed, this can be also seen in the simulation results. Not only do the proportions converge to steady states, they do this very quickly regardless of the initial conditions. This result supports the experimental setup: 1) the proportions stabilize much earlier than the detection time, so regardless of the stage at which the tumour is discovered and resected, the QAD proportions capture its intrinsic properties; 2) when performing PDX experiments, even if the piece of tumour sample used was not representative of the original patient QAD proportions at the time of injection (initial conditions), the system quickly stabilizes to those proportions governed by the parameters intrinsic to the patient.

#### 3 Parameter estimation

In order to see how our model is able to describe the dynamics observed in data, we performed parameter estimation. The proliferation rates $p_1 = p_2 = 0.55$ were kept constant, assuming a cell cycle length of 30h for GBM cells. For simplicity, $\delta = 0.2$ was also kept constant to a value informed by healthy V-SVZ, which nevertheless does not have a huge impact on the dynamics. As a result, the parameters estimated comprise of $r, b_1$ and $b_2$. The optimization was based on least squares. A cost function was defined as the sum of squared differences between the proportions $Q_p, A_p, D_p$ given by the ODE system at the end of the time interval considered, and the QAD proportions obtained with *ptalign* for the 51 patient tumours. Of note, as the QAD proportions from the model very quickly converge to steady states, by simulating the tumour growth for 250 days and taking the values at the end time point is certain to be representative of the intrinsic properties (also at earlier times, for example of detection or resection). The cost function was then minimized by using the `fmincon` routine in MathWorks MATLAB software, with the sequential quadratic programming (`'sqp'`) algorithm. The algorithm was run multiple times starting from 5000 initial guesses for each of the 51 tumours, and the parameter set with the smallest cost function value out of the 5000 multi-start points was selected. Fig. shows that the mathematical model is indeed able to capture all the possible QAD compositions of tumours observed in the data.

### 4 Application of the model

As a next step, we ordered the tumours according to the "time to detection" (tD), defined as the time at which the total number of cells $Q + A_1 + A_2 + D$ reaches $10^{11}$. The "time to equilibrium" (tE), computed as the maximum time by which all of the QAD proportions in the model reach 0.99 of the respective values at the end of the time interval ($t_{end} = 250$ days), was always much smaller then the individual time to detection. The growth rate was computed by fitting an exponential function to the dynamics of the total number of cells. Additionally, we looked at the influence of the three estimated parameters on the growth rates of the tumours. The simulation results suggest that the activation rate $r$ represents a first predictive layer of GBM progression, and we notice three separate regimes. In the small and the large regimes, the activation rate is truly predictive of prognosis, whereas in the middle range, inspecting the self-renewal $b_1$ as an additional layer is needed. This supports the biological insights that the transitions between $Q$ and $A$ are the most important point on the pseudotime, and it is here where interventions could have the greatest impact.

#### References

[1] Georg Frobenius et al. *Über matrizen aus positiven elementen*. Reichsdr., 1908.

[2] Chi-Kwong Li and Hans Schneider. "Applications of Perron–Frobenius theory to population dynamics". In: *Journal of mathematical biology* 44.5 (2002), pp. 450–462.

[3] Frederik Ziebell, Ana Martin-Villalba, and Anna Marciniak-Czochra. "Mathematical modelling of adult hippocampal neurogenesis: effects of altered stem cell dynamics on cell counts and bromodeoxyuridine-labelled cells". In: *Journal of The Royal Society Interface* 11.94 (2014), p. 20140144.

[4] Frederik Ziebell et al. "Revealing age-related changes of adult hippocampal neurogenesis using mathematical models". In: *Development* 145.1 (2018), dev153544.

[5] Georgios Kalamakis et al. "Quiescence modulates stem cell maintenance and regenerative capacity in the aging brain". In: *Cell* 176.6 (2019), pp. 1407–1419.

[6] Diana-Patricia Danciu et al. "Mathematics of neural stem cells: Linking data and processes". In: *Cells & Development* 174 (2023), p. 203849.
